# Do Deep Learning Models for Co-Folding Learn the Physics of Protein-Ligand Interactions?

**DOI:** 10.1101/2024.06.03.597219

**Authors:** Matthew R. Masters, Amr H. Mahmoud, Markus A. Lill

## Abstract

Co-folding models are the newest innovation in deep-learning-based protein-ligand structure prediction. The recent publications of RoseTTAFold All-Atom and AlphaFold 3 have shown high-quality results on predicting the structures of proteins interacting with small-molecules, other proteins and nucleic-acids. Despite these advanced capabilities and broad potential, the current study presents critical findings that question these models’ adherence to fundamental physical principles and its susceptibility to overfitting specific protein families. Through adversarial examples based on established physical, chemical, and biological principles, we demonstrate notable discrepancies in protein-ligand structural predictions when subjected to biologically plausible perturbations. These discrepancies reveal a significant divergence from expected physical behaviors, indicating potential overfitting to particular data subsets within its training corpus. Our findings underscore the models’ limitations in generalizing effectively across diverse biological structures and highlight the necessity of integrating robust physical and chemical priors in the development of such predictive tools. The results advocate a measured reliance on deep-learning-based models for critical applications in drug discovery and protein engineering, where a deep understanding of the underlying physical and chemical properties is crucial.

## 1 Introduction

The release of AlphaFold 2 (AF2), RoseTTAFold (RF), and related models marked a seminal moment in computational biology and revolutionized protein structure prediction. AF2 and RF were able to leverage evolutionary sequence and structural template data to create a powerful deep learning model, capable of predicting proteins with breakthrough accuracy [1, 2]. The impact of these models has been profound, spawning hundreds of subsequent papers including high-throughput structure prediction [3, 4, 5], ensemble sampling [6, 7, 8], molecular docking [9, 10, 11, 12], stability prediction [13, 14, 15, 16], further method development [17, 18, 19, 20], and a host of other applied studies [21, 22, 23, 24].

Recently, the works of AlphaFold 3 (AF3) and RoseTTAFold All-Atom (RFAA) have extended these capabilities to a broader array of biomolecular complexes, incorporating interactions with proteins, nucleic acids, and small molecules within a single predictive framework [25]. This approach, where the protein structure is predicted simultaneously with the ligand has been coined co-folding [26]. By using a diffusion-based architecture, AF3 was able to remove many of it’s complexities such as the stereochemical loss, amino-acid specific frames, and special handling of bonding patterns. The network also de-emphasises the importance of protein evolutionary data and opts for a more generalized, atomic interaction layer. These changes allowed AF3 to train on nearly all structural data which extended it’s modelling capabilities to new tasks, such as protein-ligand and protein-nucleic acid complexes.

When benchmarked against existing molecular docking tools, RFAA and especially AF3 showed extremely high performance. In terms of blind docking of small molecules to proteins with the PoseBusterV2 dataset [27], AF3 achieved an accuracy of around 81% for predicting the native pose within 2 Å RMSD compared to the previous highest value obtained by DiffDock with 38% [28]. When the binding site is provided, traditional physics-based docking methods such as AutoDock Vina only reach an accuracy of about 60% compared to AF3 with over 93% [29]. This extremely high accuracy is likely approaching experimental-level accuracy due to the dynamic nature of small molecules, especially in solvent exposed and variable regions [30]. These groundbreaking results highlight AF3’s potential to disrupt conventional computational biology. However, they also raise questions about the physical robustness and generalization of these models in comparison to those based on physics.

In this paper, we aim to investigate the robustness of deep-learning-based co-folding models for predicting protein-ligand complexes by exposing the model to a series of adversarial examples. Our goal is to reveal the extent to which these models understand or fail to understand the underlying physical and chemical principles it is intended to simulate. By demonstrating the presence of vulnerabilities in these cutting-edge tools, we highlight the need for further improvements in the development of AI-driven structure prediction, ensuring they are not only accurate but also generalizing to unseen protein-ligand systems and reliable in their modelling of molecular physics.

## 2 Background and Related Work

The field of protein structure prediction has been revolutionized by deep learning models like AlphaFold 2 and RoseTTAFold, which significantly improved accuracy in predicting protein structures [1, 2]. However, previous studies have shown that even small, biologically plausible perturbations can result in significant discrepancies in predicted structures, highlighting vulnerabilities in these models [31]. These findings resonate with broader machine learning research that shows it is common for deep learning models to robustly interpolate even in high-dimensional cases but fail to extrapolate when challenged by novel unseen inputs [32, 33].

Newly released versions of both AlphaFold and RoseTTAFold have moved towards diffusion-based approaches that aim to model arbitrary chemical structures under one unified framework and have shown impressive results [25, 34]. These unified methods have been compared against several previous deep learning models which aimed to solve the task of protein-ligand docking directly, such as EquiBind [35] and DiffDock [28], as well as more conventional physics-based docking engines such as AutoDock Vina [29] and Gold [36]. In addition to the targeted adversarial attacks demonstrated against AF2, deep learning models for docking have also shown to be susceptible to non-physical artifacts, such as steric clashes and stretched bonds [28, 37]. Furthermore, some studies have shown that the performance of these deep learning methods predominantly comes from their pocket finding ability and not an ability to resolve detailed molecular interactions [38, 39]. Finally, deep learning models are also utilized for the scoring of docked poses, which have also come under scrutiny for their inherent bias and inability to understand physics [40, 41].

The approach to generating adversarial attacks in prior studies relies on computationally searching for small perturbations to the input that produce a large change in the output. In order to find adversarial examples against AF2, researchers searched for mutations that lead to structures with high RMSD, while maintaining high BLOSUM similarity, helping to ensure that perturbations remain within biologically plausible limits [31]. This approach finds “accuracy cliffs” within the networks and demonstrates that they are not robust to their inputs. This flaw is present in nearly all deep neural networks without specific continuity constraints [42, 43, 44].

In this paper, we take a different approach to generating adversarial examples with a specific focus on the modeling of protein-ligand complexes. Rather than carefully crafted inputs that lead to a large output deviation, we crafted adversarial examples based on known physical, chemical, and biological first principles. These examples have greater interpretability and are able to demonstrate other issues with the model, such as overfitting and it’s lack of physical understanding.

## 3 Methodology

In this study, we perform several types of adversarial challenges by crafting examples with an expected physical outcome and seeing if AF3 predictions agree with this expectation (See Footnote ^4^). Currently, AlphaFold3 is only available as a web-server with limited small-molecule docking capabilities to test. There are a set of 19 available ligands that are endogenous and commonly occurring in the PDB, such as ATP, hemes, and fatty acids. Due to the frequent appearance of many of those ligands in the PDB, it is likely that AF3 has seen most, if not all, of their possible binding pockets. Therefore, in order to assess the robustness of small-molecule predictions, we are limited to these ligands and small peptides.

### 3.1 Binding Site Mutagenesis

#### Removing Interacting Residues

In this challenge, we identify residue side-chains that interact favorably with a ligand and mutate them into glycine residues, effectively removing the driving force for binding. For this example, we selected several systems which bind one of the available ligands: FtsE which binds ATP (PDB: 8X61), CYP109B4 which binds heme (PDB: 7Y97), and lipid transfer protein which binds palmitic acid (PDB: 1MZM). These represent several diverse test cases for ligand binding that cover a number of different properties including charge, flexibility, and hydrophobicity. These targets were randomly selected and there was no automated search for complexes that fail. Supplying the unperturbed sequence as input to AF3 in blind-docking mode, AF3 produces good results with the ligand placement in agreement with the crystal structures (Figure 1). Residues forming contacts with the ligand (<3.5 Å) were selected to be mutated to glycine. For the ATP-binder, these residues were: Y11, R15, S37, K41, S42, T43, Q86, K130, S139, E142, Q163, and H195. For the heme-binder, these residues were: N69, M84, H92, R96, T246, L250, I292, R294, H350, and C352. For the fatty acid binder, these residues were R46, A57, P80, Y81, and I83. Following mutagenesis of the sequence, structures were generated again using AF3, with the highest confidence model selected. RMSD of ligand heavy atoms to the native crystal pose were computed for each generated structure.

**Figure 1.**
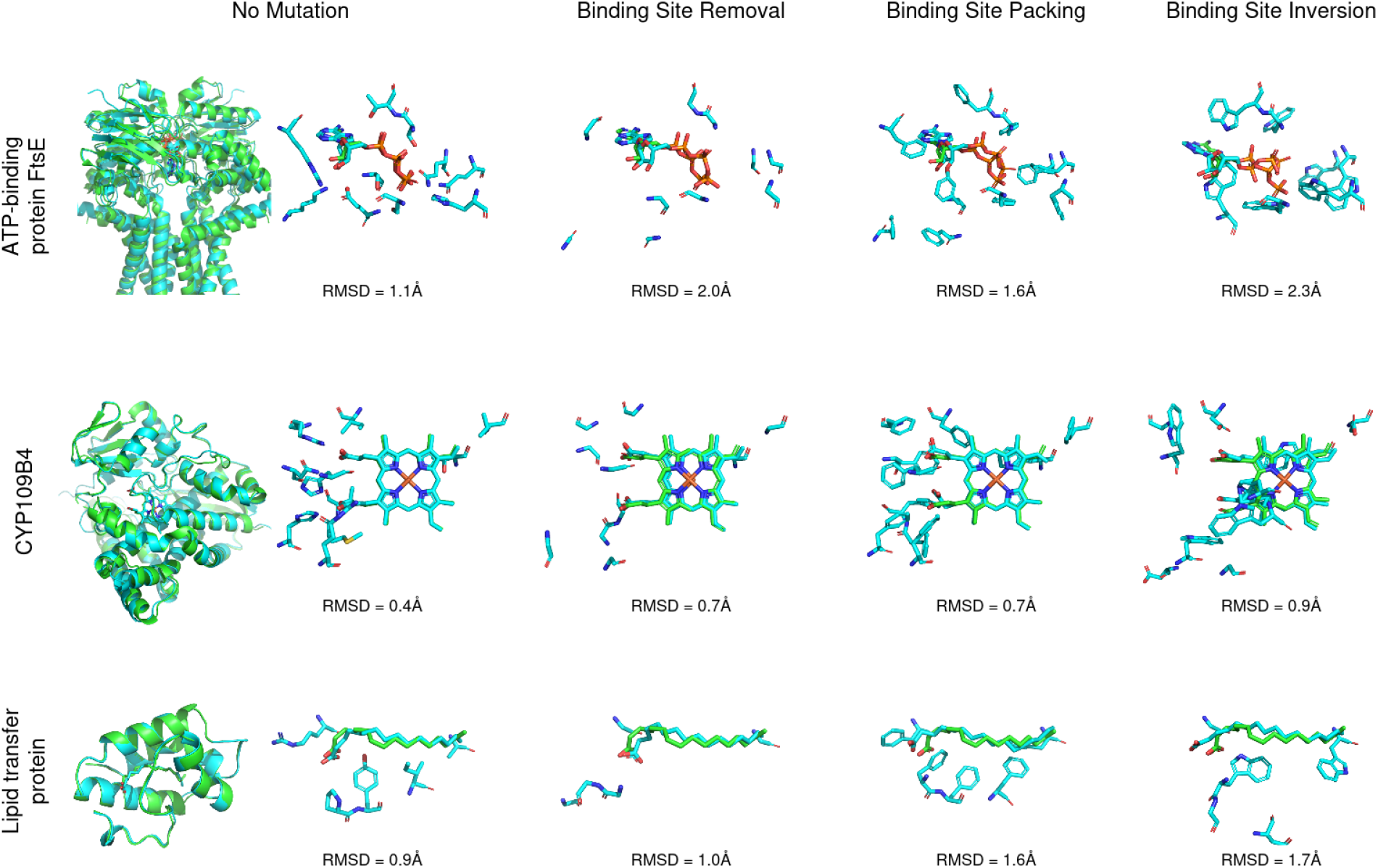
Adversarial challenges on AF3’s capacity to predict protein-ligand complexes by destruction of the nature of the binding sites. Studied systems displayed are ATP-binding protein FtsE, heme-containing CYP109B4, and lipid transfer protein. All binding site residues contacting the ligand (<3.5 Å) were mutated to glycine (removal of interactions with side chains), phenylalanine (packing of binding site), and to dissimilar residues (inversion of binding site). These mutations should annihilate the binding site and remove the majority of native protein-ligands interactions necessary for binding. However, in all cases the ligand is predicted with a near identical pose, indicating that AF3 is not predicting poses based on physics of interactions, but rather learning patterns in the global protein structure and sequence.

#### Packing with Bulky Hydrophobes

In addition to the previous approach where we replace critical residues with glycine, we also mutated the same residues with phenylalanine residues. This task not only destroys the interactions that drive ligand binding, but also occupies the binding site with bulky, hydrophobic groups. Based on physico-chemical knowledge, we would expect these groups to avoid contact with the solvent, and especially highly polar groups like the triphosphate of the ATP. The aim of this challenge is to mimic the hydrophobic effect and provide an additional penalty to try and displace the ligands from their binding pockets. We used the same systems and mutation locations as the previous approach (Figure 1).

#### Mutating to Dissimilar Residues

In a final binding site mutagenesis challenge, we chose to mutate residues into those with opposing properties. For example, a residue with a small, polar side-chain (e.g. serine) is replaced by a large, non-polar residue (e.g. tryptophan), and vice-versa. This challenge goes one step further, not only removing all interactions or crowding the binding site, but actually replacing favorable interactions with unfavorable ones. To this end, we utilized the Miyata distance which gives a quantitative measure of how similar two amino acids are based on two properties: volume and polarity [45]. Again, the same three systems introduced earlier were used in this challenge. The mutations applied to each binding site are described in the Supplementary Information Section A.1.

### 3.2 Peptide Binder Mutagenesis

In addition to the select number of ligands available to dock with AF3, there is also the ability to model peptides with a minimum length of four. These tetramers are similar in size and properties to standard small-molecule drug-like molecules, so they make an apt comparison in lieu of real drug molecules. Therefore, we also selected several small peptide docking examples in order to test mutations on the ligand side as well.

The first system is the TonB-dependent receptor YncD (PDB: 6V81) which features a binding site with large positive charge that is known to bind quinoline in it’s charged state of -3 [46, 47] (Figure 2A). It is reasonable to expect that this binding site can accommodate a small, negatively charged peptide. To test this hypothesis, we predicted the receptors structure in complex with the negatively charged aspartate tetrapeptide, DDDD (Figure 2B). Unsurprisingly, it was able to place the tetrapeptide into the positively charged binding site. The prediction was repeated with tetrapeptide FFFF, a non-polar peptide unlikely to bind into such a charged pocket (Figure 2C).

**Figure 2.**
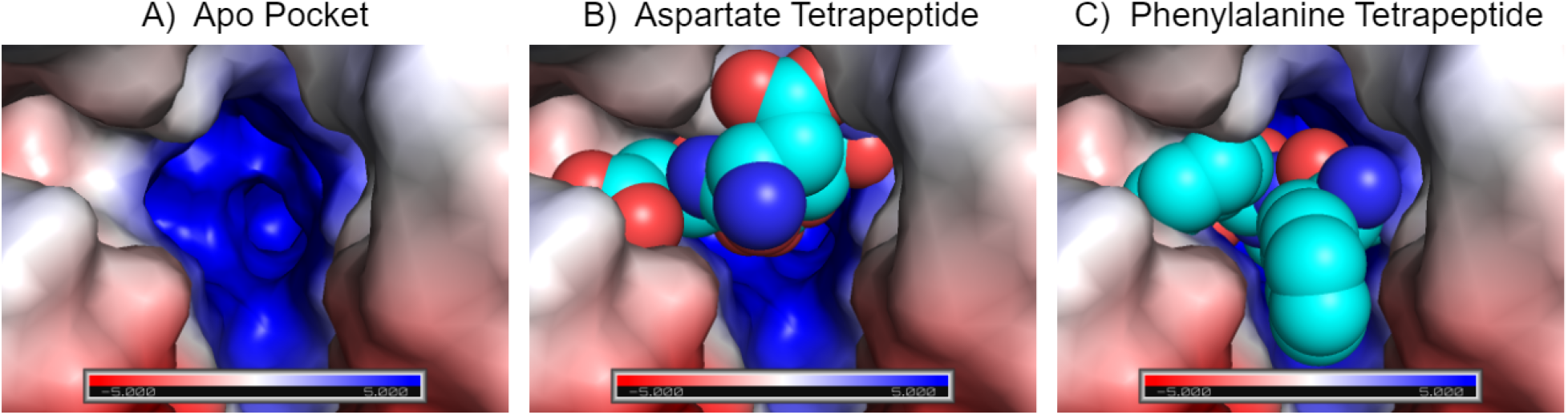
Polar vs non-polar tetrapeptide binding challenge. A) YncD features a strongly positive binding pocket that binds a cofactor with three negatively charged carboxyls. B) Aspartate tetrapeptide binds to positively charged pocket as expected. C) Non-polar phenylalanine tetrapeptide which is unlikely to bind in such a charged pocket, yet AF3 still places it inside.

The other systems include the mu-opioid receptor, which has a known high-affinity tetrapeptide binder called endomorphin (PDB: 8F7R) [48], sortase A which binds a pentapeptide (PDB: 1T2W), and gamma chymotrpsin which also binds a pentapeptide (PDB: 1GMD). The aim of this challenge was intended to probe the models behavior under mutations on the ligand side, similar to what was done in Section 3.1. However, for all three of these small peptide test systems, AF3 failed to produce the correct pose even in the unmutated case, with *C*_*α*_ RMSD values of 4.5 Å, 4.9 Å, and 4.7 Å for the three systems respectively. Due to these failures, we were unable to create additional mutation challenges based on these systems. Predicted poses for these test systems can be seen in the Supplementary Information Figure S1.

## 4 Results and Discussion

The results of the binding site mutagenesis challenges are shown in Figure 1. In the first system, AF3 predicted the ATP binding pose to within 1.1 Å RMSD of the crystal pose. We selected key interactions that form salt bridges and hydrogen bonds to the ribose and triphosphate moieties and pi-pi interaction with the adenine moiety of ATP, anchoring the ligand into the binding site. These residues were mutated to glycine and phenylalanine respectively for the “removal” and “packing” challenges. The first challenge is intended to remove all meaningful short-range interactions between the ligand and it’s known binding site. This mutation of 12 residues should remove any potential of the binding site to bind ATP. However, AF3 predicts nearly the same pose with RMSD of 2.0 Å despite having lost nearly all contacts with the protein. Therefore, we are left to deduce that AF3 predicts this pose of ATP not based on molecular interactions, but rather patterns observed in regions of the protein distant from the binding site which should play little to no role in ATP-binding, or patterns in the overall fold and sequence of the ATP binding protein.

In the next challenge, we mutate the same residues to phenylalanine, essentially packing the binding site with large, hydrophobic groups. Based on physical and chemical intuition, we would expect these phenylalanine residues to avoid contact with solvent and especially highly negatively charged groups such as the phosphate groups of ATP. Despite performing blind docking which would allow AF3 to place ATP anywhere on the surface of FtsE, AF3 disregards the physical-chemical principles of moleculsr interactions and continues to place the ATP molecule as top-pose in the same position as in the wild-type binding site (RMSD: 1.6 Å). AF3 appears to accommodate the placement of the surrounding phenylalanines in a reasonable way that mostly avoids clashes, but still disregards the need of favorable interactions to form the protein-ligand complex. This reaffirms the previous statement that AF3 is not placing the ligand based on a physically-driven interactions, but rather by non-interaction patterns it learned during training.

In the final challenge, we mutate binding site residues individually into ones with opposing properties. This challenge not only removes favorable interactions, but replaces them with unfavorable ones and significantly changes the shape of the binding pocket. Despite the drastic changes to the mutated structure, the ATP ligand remained bound. In contrast to the previous challenges, this resulted in a pose with > 2.0 Å RMSD to the native due to some conformational change within the triphosphate group. However, this change is still relatively minor and does not agree with the physically expected output.

In the heme and fatty acid binding proteins, a similar trend emerged. In the unmutated case, the heme and fatty acid were predicted with high accuracy (RMSDs: 0.4 Å and 0.9 Å respectively). Again, after the annihilation of contacting residues, AF3 places the ligands in the exact same location (RMSDs: 0.7 Å and 1.0 Å respectively). Packing the binding site with phenylalanine residues and mutating to dissimilar residues did not force the ligand out of the binding site either. Additionally, we found that in many of these cases that the pLDDT confidence score was still quite high, between 70 and 85, indicating a high level of confidence despite removing all meaningful interactions. Furthermore, many of the structures generated with mutations also led to steric clashes and atoms being placed too close together (See Supplementary Information Figure S2). However, even in these cases, high confidence values greater than 70 were still observed, demonstrated the confidence models lack of ability to detect these clashes.

In order to probe the model with adversarial examples from both the protein and ligand perspective, we chose several additional peptide mutagenesis experiments. The TonB-dependent receptor YncD was selected as the first target for it’s strongly positively charged binding site which is known to bind negatively charged ligands. Unsurprisingly, AF3 is capable of placing a negatively charged aspartate tetrapeptide into the positively charged pocket. In order to challenge the model, we introduced a bulky, non-polar, phenylalanine tetrapeptide. AF3 responds by placing the peptide into the same pocket. Given the properties of the ligand and pocket, this binding is unlikely to occur experimentally and is further evidence that AF3 does not respect molecular interactions including electrostatics.

The results clearly demonstrate that AlphaFold3 does not resemble a robust model of physics and is capable of generating non-physical structures. While the current version of AlphaFold is limited in terms of protein-ligand modelling, a number of interesting test cases can be employed that probe the models understanding of properties such as molecular size, charge, polarity, flexibility, and so on. Due to the limited release of the current version of AF3, our initial validation attempts are rather limited with respect to accessible chemical space. With the eventual release of a less-restrictive version, a more targeted assessment of small-molecule docking capabilities and limitations of AF3 need to be made. It also would be desirable in future studies to use explainable AI techniques to explore the features that AF3 actually attends to when placing the ligands in the highly mutated binding sites of the target proteins.

## 5 Conclusion

Despite the groundbreaking capabilities of AF3 in predicting complex biomolecular structures, our initial study highlights its limitations in adhering to fundamental physical principles and a tendency to overfit specific protein families. Through adversarial examples based on established physical, chemical, and biological principles, we uncovered vulnerabilities in AF3’s predictions that contradict the scientific knowledge of molecular interactions. These findings underscore the need for more rigorous validation of deep learning models for protein-ligand structure prediction such as AF3 to be able to generate trust in this new technology for critical applications such as drug design. We emphasize that the model outcomes should be critically validated on their compliance with physical and chemical principles. Future efforts should focus on refining these tools addressing biases introduced by training data to ensure broader reliability and generalization capacity towards new chemical and biological entities.

## A Supplementary Information

### A.1 Mutating to Dissimilar Residues

As described in the main text, one of the binding site mutagenesis challenges we employed was to replace contacting residues with those that have opposite properties. In this study, we utilized the Miyata distance in order to select mutations with significant dissimilarity. The Miyata distance is defined as 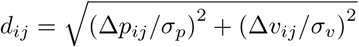 where Δ*p*_*ij*_ is difference of polarity between replaced amino acids, Δ*p*_*ij*_ is difference of volume, σ_*p*_ and σ_*v*_ are the standard deviations for Δ*p*_*ij*_ and Δ*v*_*ij*_, respectively. Therefore, here we summarize the mutations we applied during this challenge in the table below:

**Table S1:** Assignment used for mutating to dissimilar binding site residues according to the Miyata distance.

| Wild-Type | Mutation | Distance |
| --- | --- | --- |
| TYR | GLY | 4.08 |
| TRP | GLY | 5.13 |
| LYS | GLY | 3.54 |
| ARG | GLY | 3.58 |
| VAL | ASP | 3.40 |
| LEU | ASP | 4.10 |
| ILE | ASP | 3.98 |
| MET | ASP | 3.69 |
| PHE | ASP | 4.27 |
| ALA | TRP | 4.23 |
| GLY | TRP | 5.13 |
| PRO | TRP | 4.17 |
| HIS | TRP | 3.16 |
| ASP | TRP | 4.88 |
| GLU | TRP | 4.08 |
| CYS | TRP | 3.34 |
| ASN | TRP | 4.39 |
| GLN | TRP | 3.42 |
| THR | TRP | 3.50 |
| SER | TRP | 4.38 |

Therefore, for the ATP-binder the mutated residues were: Y11G, R15G, S37W, K41G, S42W, T43W, Q86W, K130G, S139W, E142W, Q163W, and H195W. For the heme-binder, these residues were: N69W, M84D, H92W, R96G, T246W, L250D, I292D, R294G, H350W, and C352W. For the fatty acid binder, these residues were R46G, A57W, P80W, Y81G, and I83D.

### A.2 Additional Results

#### A.2.1 Failed Peptide Examples

As noted in the main text, three different systems with a known short peptide binder were selected as additional test systems. However, when AF3 was used to predict the peptide binding mode prior to any mutation, it failed in all three cases. Therefore, we were unable to progress further and challenge the model by mutating these examples. Below we show the predicted binding mode and RMSD for each of these systems prior to any mutation:

**Figure S1:**
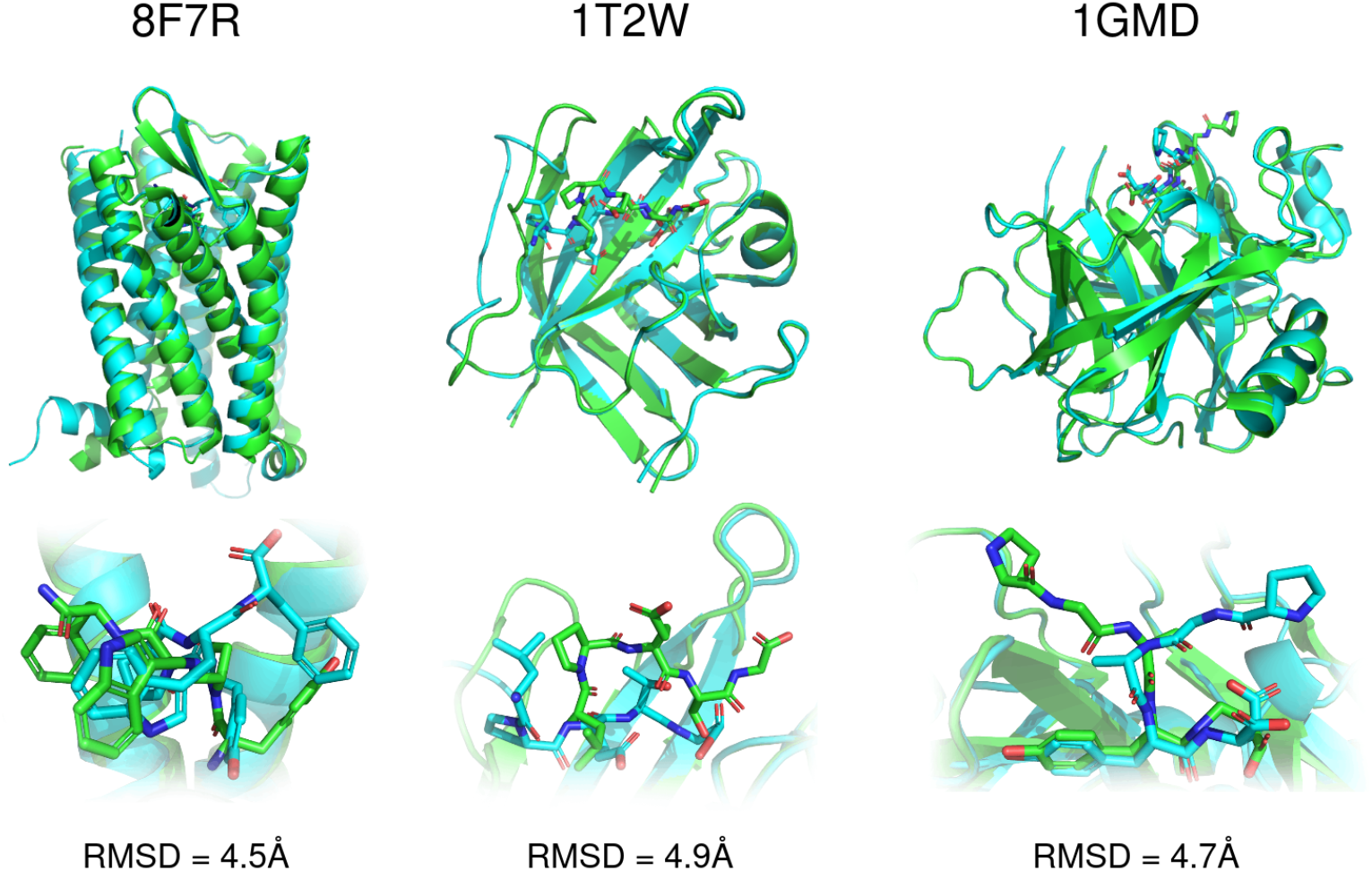
Examples of AlphaFold failure on short peptide examples without any mutation. Green is the crystal structure while cyan is the AF3 prediction. Top row shows full protein structure while bottom row focuses on the binding site.

#### A.2.2 Steric Clashes

In the mutated structures, steric clashes were often observed where heavy atoms overlapped as close as 0.4 Å. Below we show two examples of these steric clashes between the protein and ligand heavy atoms:

**Figure S2:**
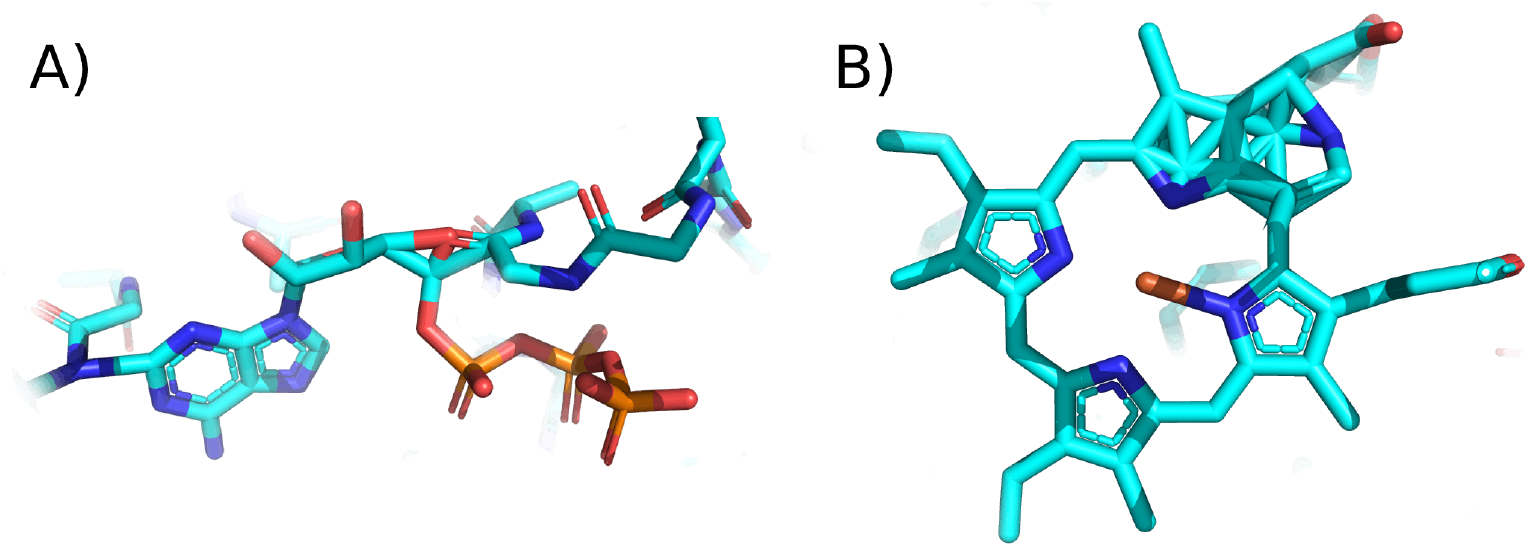
Examples of steric clashes produced in mutated structures. A) ATP example during the binding site removal challenge. B) Heme example during the binding site packing example. In both cases, atoms from the protein side-chain and backbone overlap significantly with the ligand atoms. However, the confidence scores for these ligands were still above 70, indicating a structure with high confidence.

## Footnotes

4 This study is a work-in-progress and only focuses on AF3. Further investigations into other cofolding models is planned.

